# Glucokinase activity controls peripherally-located subpopulations of β-cells that lead islet Ca^2+^ oscillations

**DOI:** 10.1101/2024.08.21.608680

**Authors:** Erli Jin, Jennifer K. Briggs, Richard K.P. Benninger, Matthew J. Merrins

**Affiliations:** Department of Medicine, Division of Endocrinology, Diabetes & Metabolism, University of Wisconsin- Madison, Madison, WI, United States; Department of Bioengineering, University of Colorado Anschutz Medical Campus, United States; Barbara Davis Center for Childhood Diabetes, University of Colorado Anschutz Medical Campus, United States

## Abstract

Oscillations in insulin secretion, driven by islet Ca^2+^ waves, are crucial for glycemic control. Prior studies, performed with single-plane imaging, suggest that subpopulations of electrically coupled β-cells have privileged roles in leading and coordinating the propagation of Ca^2+^ waves. Here, we used 3D light- sheet imaging to analyze the location and Ca^2+^ activity of single β-cells within the entire islet at >2 Hz. In contrast with single-plane studies, 3D network analysis indicates that the most highly synchronized β-cells are located at the islet center, and remain regionally but not cellularly stable between oscillations. This subpopulation, which includes ‘hub cells’, is insensitive to changes in fuel metabolism induced by glucokinase and pyruvate kinase activation. β-cells that initiate the Ca^2+^ wave (‘leaders’) are located at the islet periphery, and strikingly, change their identity over time via rotations in the wave axis. Glucokinase activation, which increased oscillation period, reinforced leader cells and stabilized the wave axis. Pyruvate kinase activation, despite increasing oscillation frequency, had no effect on leader cells, indicating the wave origin is patterned by fuel input. These findings emphasize the stochastic nature of the β-cell subpopulations that control Ca^2+^ oscillations and identify a role for glucokinase in spatially patterning ‘leader’ β-cells.

**Highlights:** • Studies of islet Ca^2+^ oscillations by 3D light-sheet imaging provide a more complete picture of β- cell subpopulations than prior 2D studies.

• Highly synchronized β-cells (including ‘hub cells’) are a regionally-stable subpopulation located at the islet center that is insensitive to metabolic perturbation.

• Glucokinase activation patterns the Ca^2+^ wave axis, which originates from stochastic β-cell subpopulations on the islet periphery that change between oscillations.

• The stochasticity of ‘leader’ β-cells, and the stability of ‘hubs’, is geographically consistent with the peripheral location of α/δ-cells in mouse islets.

## Introduction

Pulsatile insulin secretion from pancreatic islet β-cells is key to maintaining glycemic control. Insulin secretory oscillations increase the efficiency of hepatic insulin signaling and are disrupted in individuals with obesity and diabetes^1^. The primary stimulus for insulin release is glucose, which is intracellularly metabolized to generate a rise in ATP/ADP ratio, which closes ATP-sensitive K^+^ channels (K_ATP_ channels) to initiate Ca^2+^ influx and insulin secretion^2^. At elevated glucose, β-cells oscillate between electrically silent and electrically active phases with a period of minutes, both in vivo and in isolated islets. The depolarizing current can be transmitted between β-cells across the whole islet through gap junction channels. However, the activity of individual electrically coupled β-cells is functionally heterogenous^3–9^. This heterogeneity results in the emergence of β-cell subpopulations that may be crucial for maintaining the coordination of the whole islet and regulating pulsatile insulin release^10–13^. Understanding the underpinnings of β-cell functional heterogeneity and islet cell communication is important for understanding islet dysfunction and the pathogenesis of diabetes.

Similar to studies of neuronal networks, functional network analysis can be used to quantify interactions within the heterogenous β-cell system. Interactions (termed ‘edges’) are drawn between β- cell pairs with highly correlated Ca^2+^ dynamics. Studies suggest that the β-cell functional network exhibits high clustering or ‘small-world’ properties^14^, with a subpopulation of β-cells that are highly synchronized to other cells (‘hub cells’)^10^. Silencing the electrical activity of these hub cells with optogenetics was found to abolish the coordination within that plane of the islet^10,15,16^. Similarly, time series-based lagged cross correlation analysis has identified subpopulations of cells at the wave origin, termed ‘early-phase’ or ‘leader cells’, that lead the second-phase Ca^2+^ wave by depolarizing and repolarizing first^12,13^. However, questions have been raised whether the highly networked or leader subpopulations have the power to control the entire islet^15,17–19^. Underlying this controversy lies several unanswered questions: what mechanisms drive the existence of these functional subpopulations? Do these subpopulations arise primarily from mechanisms intrinsic to β-cells, making the subpopulations consistent over time? Alternatively, do they arise from the combination of intrinsic mechanisms and emergence due to surrounding cells, allowing the subpopulations to fluidly change over time? To date, experiments have been restricted to imaging a single two-dimensional (2D) plane of the islet which contains only a small fraction of the β-cells present in the three-dimensional (3D) islet tissue, limiting the ability to address these questions.

With these caveats in mind, prior studies using a mixture of computational and molecular approaches suggested that β-cell subpopulations are patterned by glucokinase, which is often referred to as the ‘glucose sensor’ for the β-cell^13,20,21^. By phosphorylating glucose in the first step of glycolysis, glucokinase activation lengthens the active phase of Ca^2+^ oscillations by committing more glucose carbons to glycolysis^22^. Until recently, it was believed that downstream glycolysis was irrelevant to pulsatile insulin secretion. However, in conflict with this model, allosteric activation of pyruvate kinase accelerates Ca^2+^ oscillations and increases insulin secretion^22,23^. As a potential mechanistic explanation for these observations, plasma membrane-associated glycolytic enzymes, including glucokinase and pyruvate kinase, have been demonstrated to regulate K_ATP_ channels via the ATP/ADP ratio^24^. However, it remains unknown whether these glycolytic enzymes influence β-cell heterogeneity and network activity.

To study single β-cell activity within intact islets, we engineered a 3D light-sheet microscope to simultaneously record the location and Ca^2+^ activity of single β-cells over the entire islet during glucose-stimulated oscillations. In concert, we developed 3D analyses to investigate the spatial features of subpopulations that underlie the β-cell network and Ca^2+^ wave, and the consistency of these features over time. We further examined the consequences of sampling islet heterogeneity in 2D compared to 3D. Finally, we investigated the role of the glycolytic enzymes glucokinase and pyruvate kinase in controlling β-cell subpopulations during glucose-stimulated oscillations.

## Results

### Light-sheet microscopy enables high-speed 3D imaging of oscillations in single β-cells within intact islets

To acquire high-speed 3D time course imaging of β-cell Ca^2+^ oscillations within intact islets, we utilized a lateral-interference tilted excitation light-sheet system^25^ mounted on an inverted fluorescence microscope (Fig. 1A and *Methods*). To image Ca^2+^ activity and spatially resolve individual β-cells in intact islets, islets were isolated from *Ins1-Cre:Rosa26^GCaMP6s/H2B-mCherry^*mice that express cytosolic GCaMP6s Ca^2+^ biosensors and nuclear H2B-mCherry reporters selectively within β-cells. We first compared the images collected by the light-sheet system with a commercial spinning disk confocal using the same 40× water immersion objective. Similar to a widefield microscope, the axial resolution of the light-sheet microscope is dictated by the numerical aperture (NA) of the objective lens (∼1.1 μm for a 1.15 NA objective and GCaMP6s emission)^25^, whereas the spinning disk uses a pinhole array to enhance axial resolution. At a shallow depth of 24 µm from the coverslip, H2B-mCherry-labelled nuclei and GCaMP6s-labelled β-cells were resolved both by the light-sheet and the spinning disk confocal. However, the nuclei were only resolved by the light-sheet system at depths ≥ 60 μm due to the reduced light scatter from side illumination (Fig. 1B). Thus, the main advantage of the light-sheet system is the ability to image the entire islet in 3D (Fig. 1C).

**Fig. 1.**
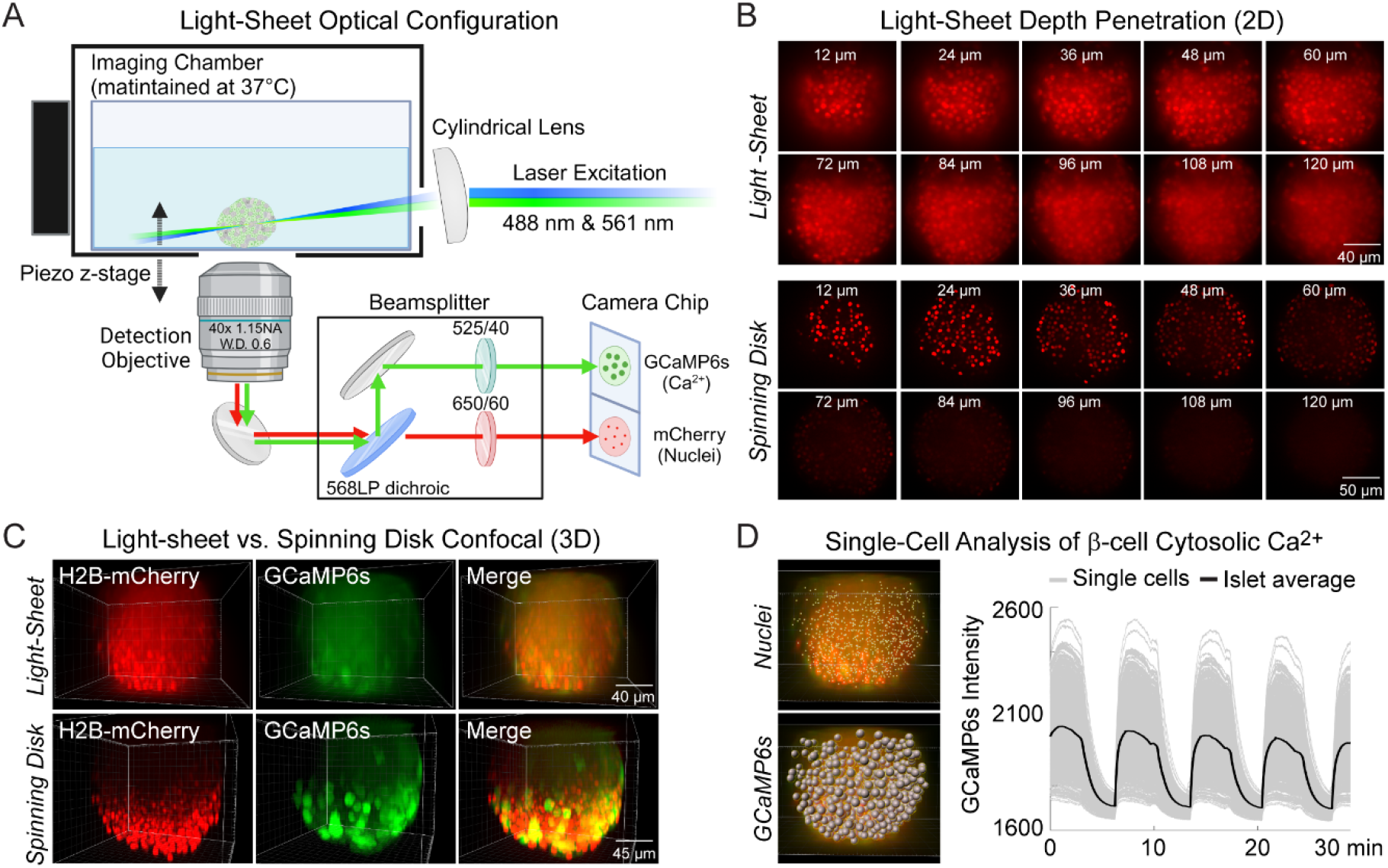
Engineering of a light-sheet microscope to image intact islets in 3D. (A) Schematic of the light-sheet microscope showing the optical configuration. (B) Representative light-sheet (upper panel) and spinning disk confocal images (lower panel) of a mouse pancreatic islet expressing β-cell specific H2B-mCherry fluorophore at different 2D focal planes, emphasizing the superior depth penetration of the light-sheet microscope. (C) 3D imaging of β-cells expressing GCaMP6s Ca^2+^ biosensors and nuclei mCherry biosensors. (D) Using *Ins1-Cre:Rosa26^GCaMP6s/H2B-mCherry^* islets, the software-identified center of β-cell nuclei (yellow dots) was used to generate GCaMP6s regions of interest (gray spheres). A representative Ca^2+^ timecourse is displayed in the right panel for an islet stimulated with glucose and amino acids.

Prior studies of β-cell Ca^2+^ oscillations utilized 1 Hz imaging to resolve phase shifts for single β- cell traces within a single 2D plane^26–29^. To image the entire islet at similar acquisition speeds, the hardware was operated under triggering mode to minimize communication delays (Suppl. Fig. 1). In this mode, the lasers and the piezo z-stage were triggered directly by the camera, which received a single set of instructions from the computer via the NiDAQ card. To image 132 µm into the islet at 2 Hz, 15 ms was allowed for photon collection and stage movement for each of the 4-µm z-steps. Initially, an hour-long delay was required to save 120,000 imaging files after running a continuous 30-minute experiment. Because this delay is only observed after the first 3 minutes of imaging, it was possible to eliminate the delay by separating the acquisition into a series of 3-minute loops (Suppl. Fig. 2).

Individual β-cell nuclei were located using the Spots function of Bitplane Imaris software (Fig. 1D). For each nucleus, a cellular region of interest (ROI) was defined by a sphere of radius 4.65 microns around the ROI center based on results from a computational automated radius detection^30^. The mean GCaMP6s intensity of each cell ROI for each time point were calculated and exported as single cell traces. We did not observe any significant photobleaching using continuous GCaMP6s and H2B- mCherry excitation over the course of the experiment. The combination of this light-sheet system, islet cell labeling, and analysis pipeline allows for imaging of Ca^2+^ from nearly all β-cells in the islet at speeds fast enough for spatio-temporal analyses to identify functionally heterogeneous β-cell subpopulations.

### 3D analyses of islet Ca^2+^ oscillations reveal that the β-cell network is distributed in a radial pattern while Ca^2+^ waves begin and end on the islet periphery

To investigate the synchronization between β-cells across the islet in 3D space, we imaged and extracted Ca^2+^ time-courses for *Ins1-Cre:ROSA26^GCaMP6s/H2B-mCherry^*islets that exhibit slow oscillations. Following the network analysis methods set forth in^14,31^, we calculated the correlation coefficient between every cell pair and defined an “edge” between any cell pairs whose correlation coefficient was above threshold (Fig. 2A). This threshold was set such that the average number of edges per cell, also called the ‘cell degree’, was equal to 7. A fixed average degree rather than fixed threshold was used to mitigate inter-islet heterogeneity^31^. An example 3D network for a single β-cell within an islet is shown (Fig. 2B) along with the frequency distribution of all β-cells within the islet (Fig. 2C). The high degree cells (top 10% of the total population, *blue*) and low degree cells (bottom 10% of the total population, *red*) were then mapped onto a 3D projection of the islet and onto the Ca^2+^ time course (Fig. 2D).

**Fig. 2.**
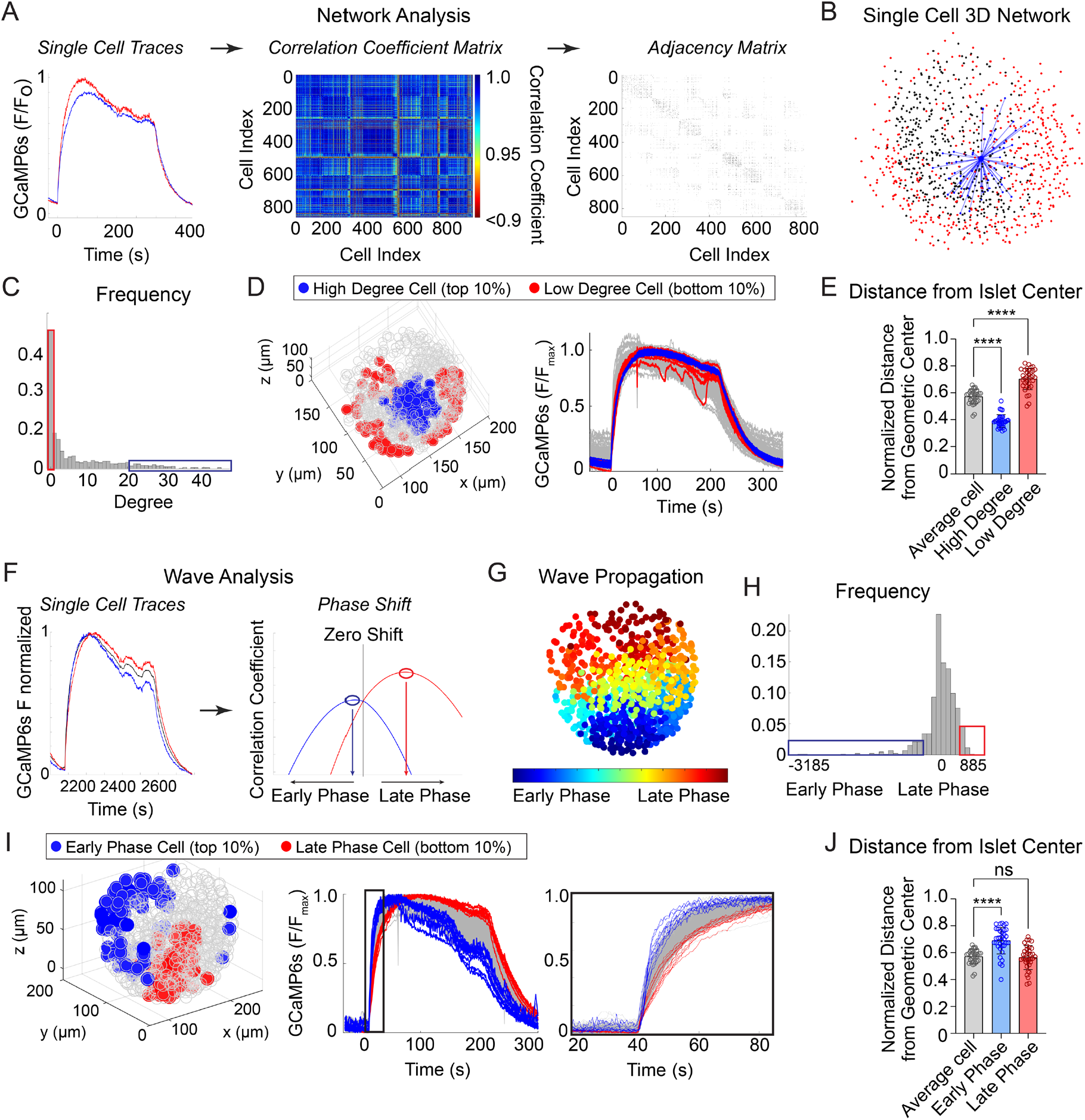
Characterization of single β-cells using 3D network and phase analysis. (A) Flow diagram illustrating the calculation of cell degree from pairwise comparisons between single β-cells. (B) An example 3D network for a single β-cell within a representative islet is shown with synchronized cell pairs in blue, cells that have other synchronized pairs in black, and cells that are asynchronous in red. This analysis is repeated for all cells in the islet. (C) Frequency distribution of cell degree for all β-cells analyzed. Top 10% (blue box) and bottom 10% (red box) are high and low degree cells. (D) Representative 3D illustration and Ca^2+^ traces showing the location of high degree cells (blue) and low degree cells (red). (E) Quantification of the normalized distance from the islet center for average degree cells (gray), high degree cells (blue), and low degree cells (red). (F) Flow diagram illustrating the calculation of cell phase, calculated from the correlation coefficient and phase shift. (G) Wave propagation from early phase cells (blue) to late phase cells (red) in 3D space. (H) Frequency distribution of cell phase for all β-cells analyzed. Top 10% (blue box) and bottom 10% (red box) are early and late phase cells. (I) Representative 3D illustration and Ca^2+^ traces showing the location and traces of high phase cells (blue) and low degree cells (red). (J) Quantification of the normalized distance from the islet center for average phase cells (gray), early phase cells (blue), and late phase cells (red). Data represents *n* = 28,855 cells, 33 islets, 7 mice. Data are displayed as mean ± SEM. \*\*\*\**P* < 0.0001 by 1-way ANOVA.

Compared to average degree cells, the high degree cells were consistently located at the center of the islet while the low degree cells were located on the periphery, indicating that the islet network is distributed in a radial pattern (Fig. 2D,E).

To analyze the propagation and spatial orientation of Ca^2+^ wave in 3D space, we calculated the lagged correlation coefficient between every β-cell and the islet average, and identified the phase lag with maximum correlation (Fig. 2F). The spatial distribution of phases of an example islet is shown (Fig. 2G), along with the frequency distribution of all β-cells within the islet (Fig. 2H). The early phase cells (top 10% of the total population that depolarize first and repolarize first, *blue*) and late phase cells (bottom 10% of the total population that depolarize last and repolarize last, *red*) were then mapped onto a 3D projection of the islet and onto the Ca^2+^ time course (Fig. 2I). Unlike the islet network, for which the high degree cells emanate from the islet center (Fig. 2D,E), the early phase and late phase cells were each located at the islet periphery, and show a clear temporal separation between depolarization and repolarization (Fig. 2I,J).

### The location of the β-cell network is stable over time while the wave progression varies

We next assessed the stability of high degree cells and early phase cells over time, by assessing their presence across consecutive oscillations (Fig. 3A,B; Suppl. Fig. 3). The high degree cells and early phase cells have a similar ∼60% retention rate between oscillations (Fig. 3C). Strikingly, when we examined the center of gravity for each β-cell subpopulation, we found that the center of gravity of the early phase cells moved significantly more than that of the high degree cells (Fig. 3D). This indicates that the early phase cells tend to change their identity more with each oscillation. To further investigate the change in location of early phase cells, we used principal component analysis to identify the principal axis between early phase cells and late phase cells (wave axis) and calculated the rotation of the axis between each oscillation. Of the 25 islets examined, 57% show substantial changes in the wave axis over time (Fig. 3E). Thus, β-cell depolarization is initiated at different locations within the islet over time, while the β-cell network location is relatively stable.

**Fig. 3.**
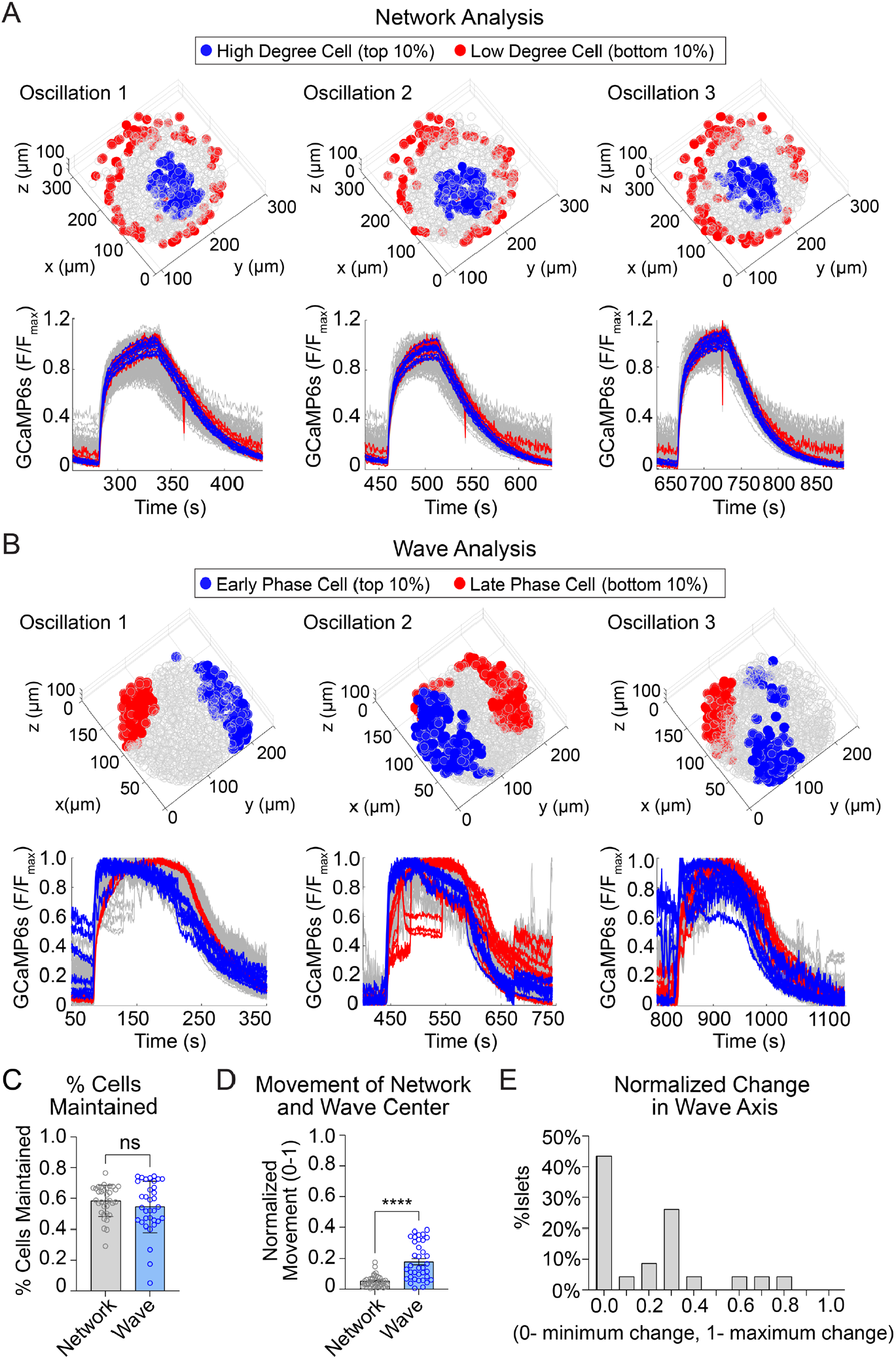
The network of highly synchronized β-cells is consistent between oscillations, while the Ca^2+^ wave axis rotates. (A) 3D representation of the islet showing the location of high degree cells (blue) and low degree cells (red) over three consecutive oscillations (top panel) and their corresponding Ca^2+^ traces (bottom panel). (B) 3D representation of the islet showing the location of early phase cells (blue) and late phase cells (red) over three consecutive oscillations (top panel) and their corresponding Ca^2+^ traces (bottom panel). (C) Quantification of the retention rate of high degree and early phase cells. (D) Relative spatial change in the center of gravity of β-cell network vs. the β-cell Ca^2+^ wave. (E) Frequency distribution showing the normalized change in Ca^2+^ wave axis for all islets. Data are displayed as mean ± SEM. \*\*\*\**P* < 0.0001 by normality test followed by Paired Student’s t-test or Wilcoxon Signed-Rank Test.

The analyses in Fig. 3 are focused on the top and bottom 10% of the population. To understand the stability of all β-cells within the 3D network or the 3D wave propagation over time, we ranked every β-cell in the islet by their phase/degree (‘cellular consistency’), as well as the spatial proximity of every β-cell to the center of gravity of the top 10% of the subpopulation (‘regional consistency’) (Fig. 4A). We quantified the change in these distributions using a normalized non-parametric, information-theoretic metric termed Kullback-Leibler (KL) divergence (see *Methods*). If the ranking of high degree cell to low degree cell (*e.g.*, A > B > C) for the first oscillation remains the same in the second oscillation, the KL divergence will be 0, indicating the cell ranking is completely predictable between oscillations. Alternatively, if the cell ranking changes between oscillations (A > C > B), the KL divergence will be 1, indicating the cell ranking is completely random (Fig. 4B). When examining the consistency of the network, the regional stability was much higher than the cellular stability over time (Fig. 4C). In contrast, when examining the wave, the cellular stability was similar to the regional stability (Fig. 4D). This analysis of KL divergence supports the previous conclusions that the β-cell network is regionally stable, but the wave can start at different locations. Additionally, because the wave was consistent cellularly, this analysis may imply that the wave is established by cellular properties, whereas the network is emergent.

**Fig. 4.**
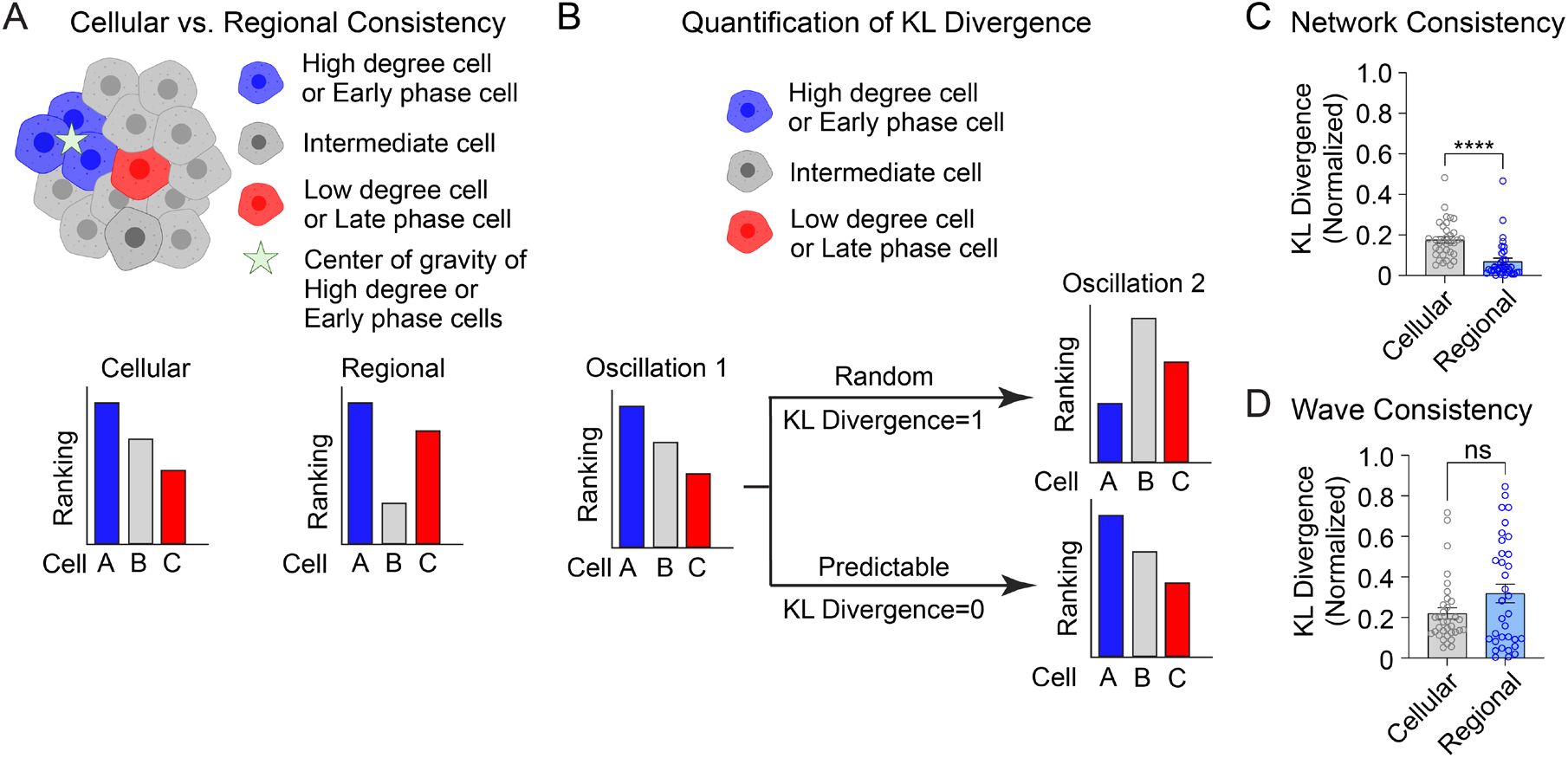
Cellular and regional consistency of the β-cell network and Ca^2+^ wave quantified by KL divergence. (A) Schematic showing cellular consistency analysis and regional consistency analysis. (B) Schematic depicting the use of KL divergence to determine consistency between consecutive oscillations. Every β-cell in the islet is ranked, with near-zero KL divergence values indicating high consistency between oscillations and near-unity KL divergence indicating randomness. (C-D) Comparison of cellular vs. regional consistency of the network (C) and wave (D) by KL divergence. Data are displayed as mean ± SEM. \*\*\*\**P* < 0.0001 by normality test followed by Paired Student’s t- test or Wilcoxon Signed-Rank Test.

### The consistency of 2D analyses of the network and wave are much lower than 3D analyses

To investigate whether 2D analysis, as performed in all prior studies^10,13,14^ provides a similar level of robustness as the current 3D analysis, we performed network and wave analyses on a single plane at either ¼-depth or ½-depth of the z-stack (Fig. 5A). Both the 2D analysis and 3D analysis showed that the wave axis changes over time (Fig. 5B). The analyses also agreed that the high degree cells are located at the center of the islet (Fig. 5C) and that the early phase cells are located at the edge of the islet (Fig. 5D). However, when we looked at the regional and cellular consistency of the β-cell network, the 2D analysis at both ¼-depth and ½-depth of the z-stack showed no difference for regional and cellular consistency (Fig. 5E). This result contradicts with the 3D analysis which showed the regional consistency of the β-cell network is significantly more stable than cellular consistency. When analyzing the wave, both 3D analysis and 2D analysis at ¼-depth showed that the cellular consistency is more stable than regional consistency, while the results from a plane at ½-depth showed no difference (Fig. 5F). These findings indicate that 2D imaging at different planes of the islet can sometimes skew the results of the heterogeneity analysis.

**Fig. 5.**
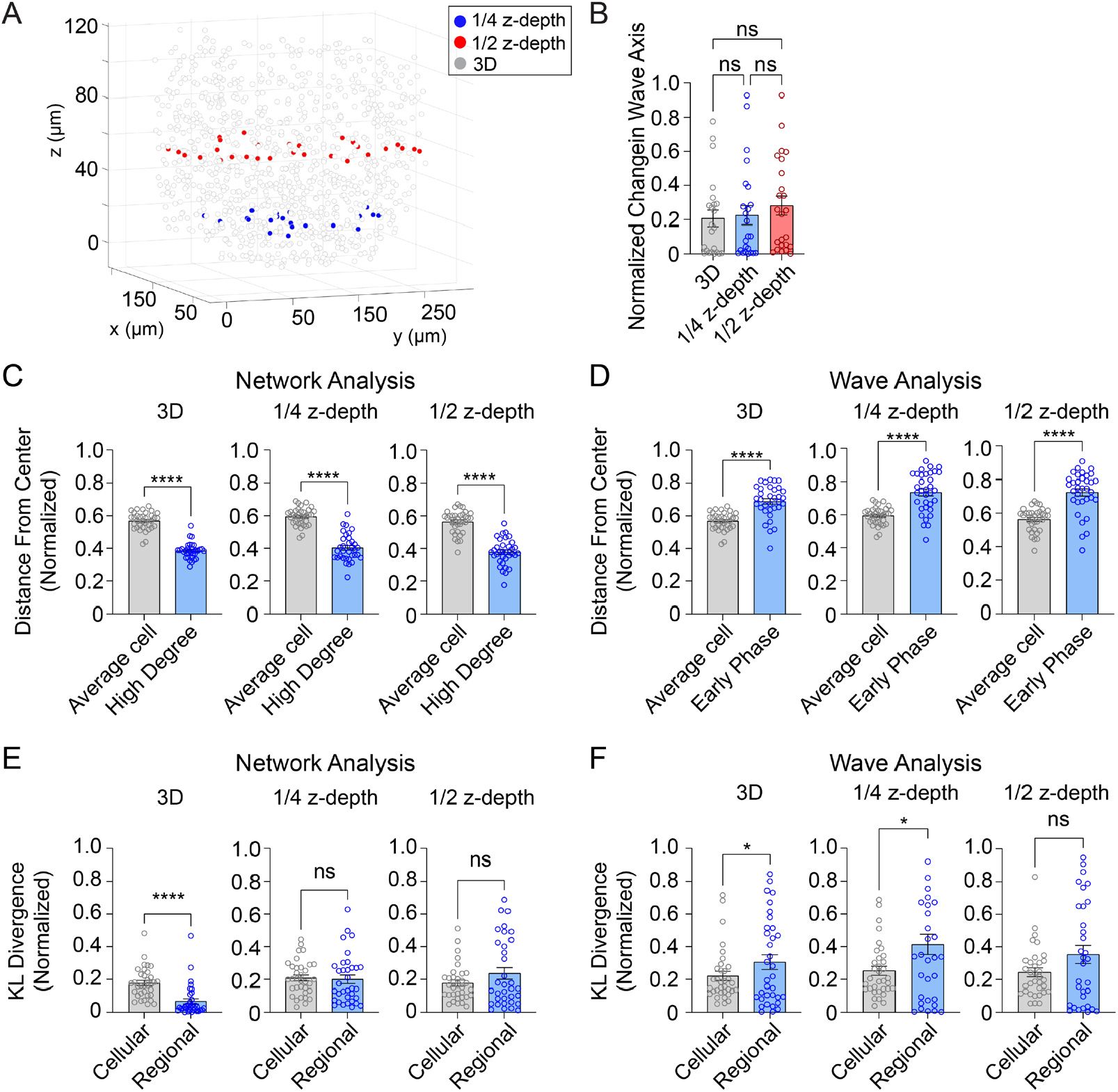
3D analysis is more robust than 2D analysis. (A) Example islet showing the locations of the ¼-depth (red) and ½-depth (blue) 2D planes used for analysis. (B) Comparison of wave axis change from 2D and 3D analysis. (C) Comparison of distance from center for average and high degree cells based on either 3D (left panel) or 2D planes (middle and right panels). (D) Comparison of distance from center for average and early phase cells based on either 3D (left panel) or 2D planes (middle and right panels). (E-F) Comparison of cellular and regional consistency of the network (E) and Ca^2+^ wave (F) based on either 3D (left panel) or 2D planes (middle and right panels). Data are displayed as mean ± SEM. \**P* < 0.05, \*\*\*\**P* < 0.0001 by normality test followed by parametric or non- parametric 1-way ANOVA (B) or Student’s t-test or Wilcoxon Signed-Rank Test (C-F).

### The origin of Ca^2+^ waves in 3D space is determined by the activity of glucokinase, while the β-cell network is patterned independently of metabolic input

Glycolysis exerts strong control over the timing of β-cell Ca^2+^ oscillations^2,32^. Glucokinase, as the ‘glucose sensor’ for the β-cell, controls the input of glucose carbons into glycolysis^2,33^, and the downstream action of pyruvate kinase controls membrane depolarization by closing K_ATP_ channels^2,22–24,34^ (Fig. 6A). We applied glucokinase activator (GKa, 50 nM RO 28-1675) and pyruvate kinase activator (PKa, 10 uM TEPP-46)^22,23^ to determine the effects of these enzymes on β-cell subpopulations during glucose-stimulated oscillations.

**Fig. 6.**
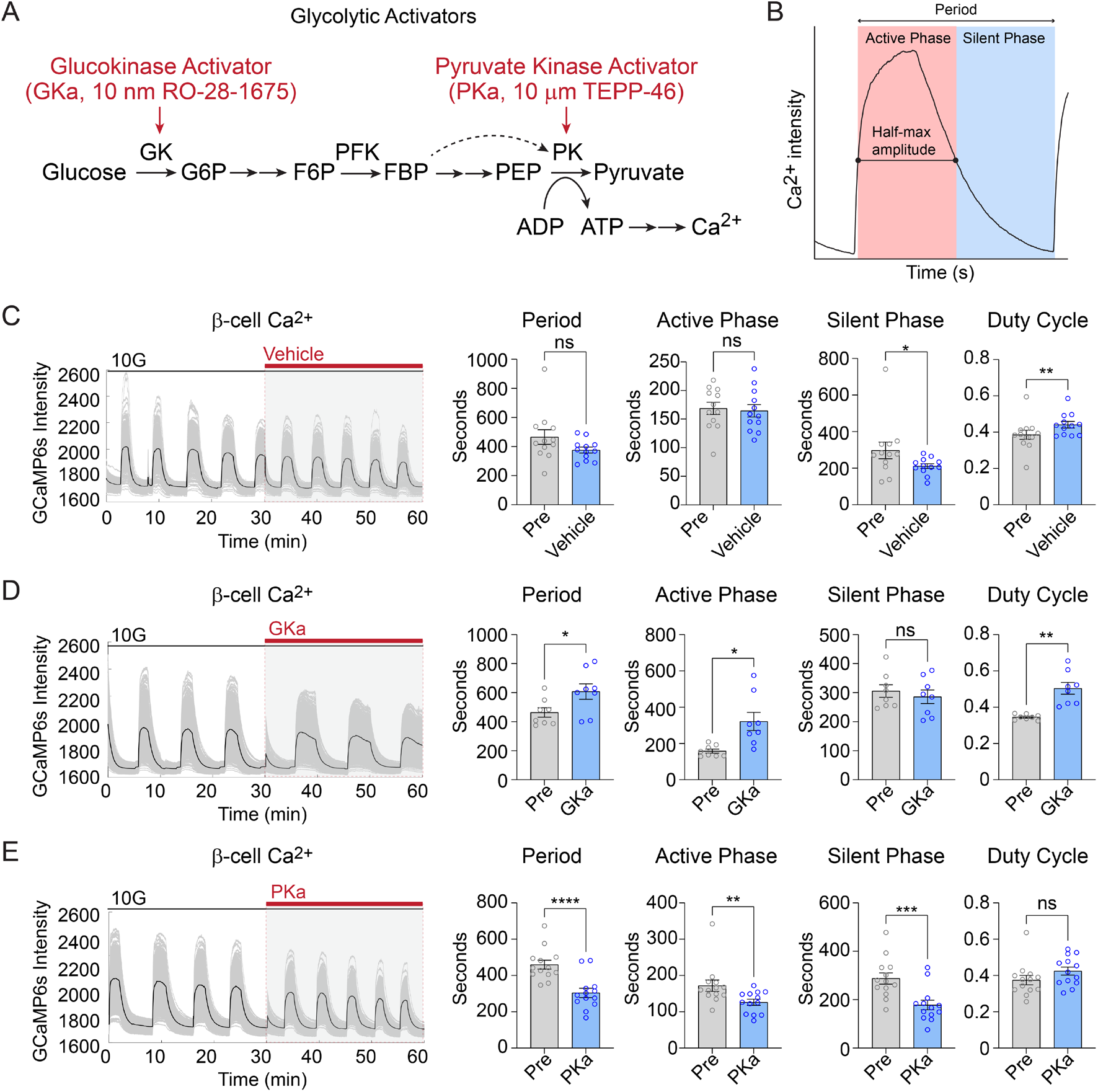
Effect of glycolytic activators on β-cell oscillations. (A) Schematic of glycolysis showing the targets of glucokinase activator (GKa) and pyruvate kinase activator (PKa). (B) Illustration indicating the oscillation period, active phase duration, silent phase duration, and duty cycle (active phase/period) calculated at half-maximal Ca^2+^. (C-E) Sample traces and comparison of period, active phase duration, silent phase duration and duty cycle before and after vehicle (0.1% DMSO) (n=11,284 cells, 13 islets, 7 mice) (C), GKa (50 nM RO-28-1675) (n=6871 cells, 8 islets, 7 mice) (D) and PKa (n=10,700 cells, 13 islets, 7 mice) (10 µM TEPP-46) (E). Data are displayed as mean ± SEM. \**P* < 0.05, \*\**P* < 0.01, \*\*\**P* < 0.001, \*\*\*\**P* < 0.0001 normality test followed by Paired Student’s t-test or Wilcoxon Signed-Rank Test.

Since biochemically distinct processes occur during the silent phase (i.e., the electrically-silent period when K_ATP_ channels close and Ca^2+^ remains low) and the active phase (i.e., the electrically-active period Ca^2+^ is elevated and secretion occurs)^2^, we quantified the duration of each phase along with the oscillation period and duty cycle (the ratio of active phase to full cycle) (Fig. 6B). In the presence of vehicle control (0.1% DMSO), the Ca^2+^ duty cycle remained stable, although an outlier in the control group resulted in a small decrease in the silent phase duration (Fig. 6C). In 3 of 15 islets, GKa induced a Ca^2+^ plateau (duty cycle = 1.0); out of necessity these islets were removed from the oscillation analysis. In the majority of islets, GKa increased the oscillation period and duty cycle (Fig. 6D). The duty cycle increase was driven by an increase in the active phase duration, with no impact on the silent phase, the time when K_ATP_ channels close (Fig. 6D). In contrast with activation of glucokinase, PKa increased the oscillation frequency by reducing the silent phase duration and the active phase duration in equal proportions (Fig. 6E). The absence of any PKa effect on the duty cycle is expected since fuel input is controlled by glucokinase, whereas silent phase shortening is expected based on the ability of PKa to reduce the time required to close K_ATP_ channels and depolarize the plasma membrane^22,23^. Thus, a single cell 3D analysis of β-cell Ca^2+^ oscillation upon GKa and PKa stimulation provides similar conclusions to prior 2D studies of intact islets.

Previous 2D studies have found metabolic differences along the Ca^2+^ wave, as measured by NAD(P)H fluorescence^13^. We measured the 3D position of early or late phase cells in response to glucokinase or pyruvate kinase activation. A positional analysis showed that GKa strongly reinforced the islet region corresponding to early phase and late phase cells, while vehicle and PKa had no discernable effect (Fig. 7A). The KL divergence for wave propagation was correspondingly reduced by GKa (Fig. 7B), indicating increased consistency, and the wave axis was significantly stabilized by GKa (Fig. 7C). Again, PKa had no discernable effect on the KL divergence for wave propagation or wave axis stability, a likely indication that the Ca^2+^ wave origin is primarily, if not exclusively, controlled by glucokinase patterning.

**Fig. 7.**
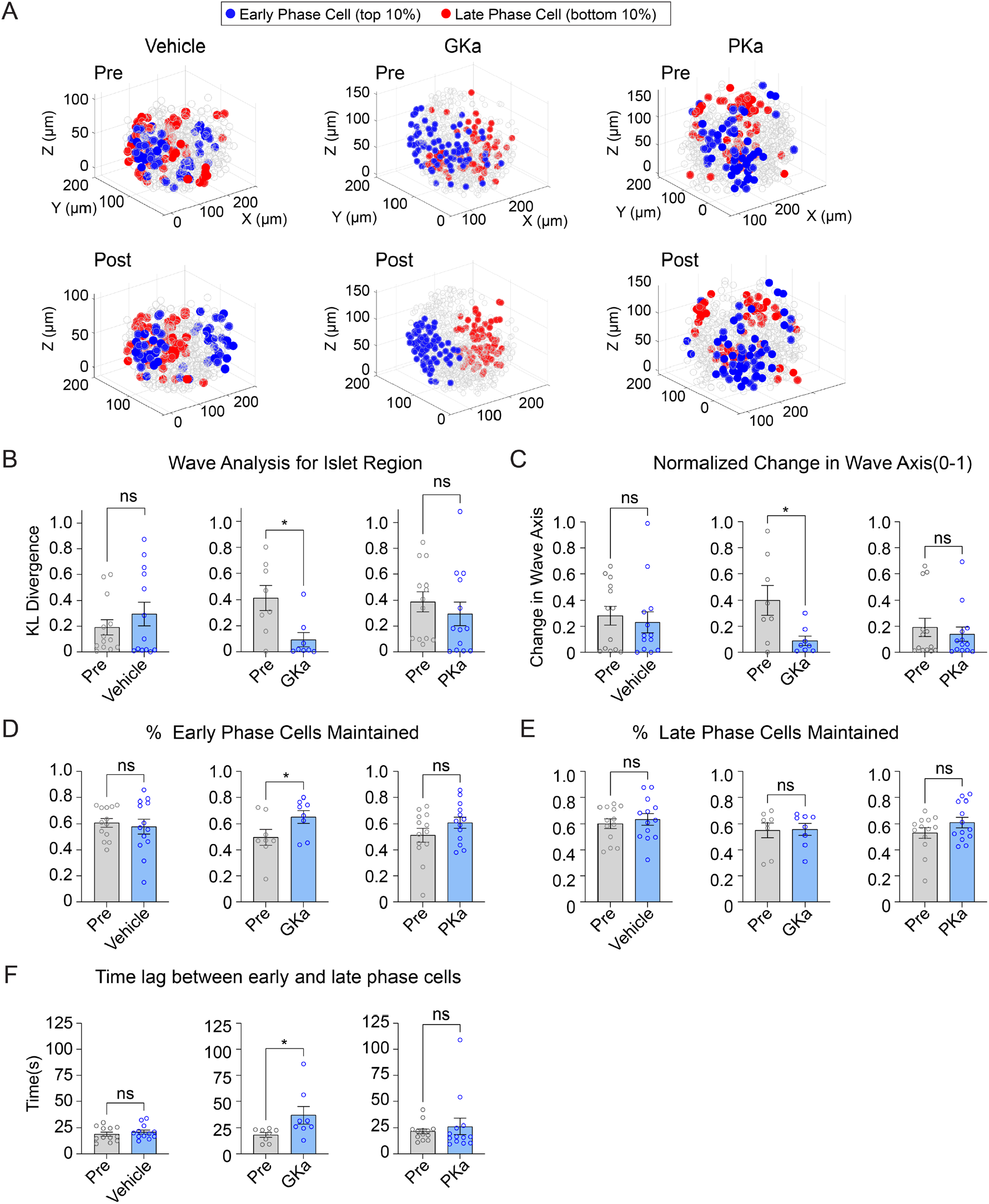
Glucokinase activity determines the origin of Ca^2+^ waves in 3D space. (A) Illustrations showing the location change of early phase cells (blue) and late phase cells (red) before and after vehicle (left panel), GKa (middle panel) and PKa (right panel). (B-H) Effect of vehicle, GKa, and PKa on regional consistency of the Ca^2+^ wave (B), wave axis change (C), early phase cell retention (D), late phase cell retention (E), and the time lag between early and late phase cells(F). Data are displayed as mean ± SEM. \**P* < 0.05 by Student’s *t*-test.

As a second approach, we examined the percentage of early phase or late phase cells maintained following activation of glucokinase or pyruvate kinase. Early phase cells were maintained to a greater degree upon GKa application, indicating greater consistency, but again showed no change upon vehicle or PKa application (Fig. 7D). Late phase cells showed no difference in their maintenance upon any of the treatments (Fig. 7E), suggesting that the earliest phase cells drive the consistency of the wave propagation. The time lag between early and late phase cells were increased upon GKa application (Fig. 7F), showing that GK activation can enlarge the differences between early and late phase cells.

While metabolic differences have been suggested to underlie functional heterogeneity in the β- cell network^10,35^, we observed no changes in the consistency of the islet network upon either GKa or PKa application (Suppl. Fig. 4A). Similarly, the consistency of high degree cells or low degree cells also did not change upon either GKa or PKa application (Suppl. Fig. 4B and 4C). Collectively, these findings implicate glucokinase as the key determinant of the Ca^2+^ wave in three-dimensional space, whereas metabolic perturbations have little influence on the islet network.

We defined early phase cells as those that depolarized and repolarized first. We also assessed whether the results were consistent for cells that only depolarized first (while ignoring repolarization). Similar to early phase cells, GKa increased the retention of β-cells that depolarized first (Suppl. Fig. 5A) and their regional consistency (Suppl. Fig. 5B). However, GKa did not influence the wave axis change (Suppl. Fig. 5C), indicating that the cells that depolarize first are a more unstable population than those that depolarize and repolarize first.

## Discussion

In this study we used light-sheet microscopy of single mouse islets to provide a three-dimensional analysis of the β-cell subpopulations that initiate Ca^2+^ oscillations and coordinate the islet network. While this *ex vivo* approach might not precisely mimic the *in vivo* situation, our analyses show that 3D imaging is a more robust approach than 2D imaging, which does not accurately reflect heterogeneity and subpopulation consistency over the entire islet. Reinforcing the concept of distinct β-cell subpopulations, the most highly synchronized cells are located at the center of the islet, while those β-cells that control the initiation and termination of Ca^2+^ waves (‘leaders’) were located on the islet periphery. We further observed that different regions of the islet initiate the Ca^2+^ wave over time, challenging the view that leader cells are a fixed pacemaker population of cells defined by their biochemistry. As discussed below, technical advances in image capture and analysis provide several new insights into the features of β-cell subpopulation in 3D and illustrate how glycolytic enzymes influence the system.

β-cell Ca^2+^ imaging is an indispensable approach for understanding pulsatility. When studied by light-sheet imaging, islets exhibited similar ∼3-5-minute oscillations as *in vivo* 2-photon imaging of β- cell Ca^2+^ oscillations in live mice^36^, as well as the high-speed confocal imaging used in prior *ex vivo* studies^37^. Light-sheet imaging overcomes the speed and depth limitations, respectively, that prevent these approaches from single-cell analysis of the entire islet. Relative to spinning disk confocal, penetration depth increased >2-fold with the light-sheet microscope (from 50-60 to 130-150 μm), allowing small and medium-sized islets to be imaged *in toto*. Abandoning confocal pinholes improved light collection, and therefore acquisition speed, ∼3-fold; this is an underestimate given the 0.4 e^-^ read noise cameras on the spinning disk microscope versus 1.6 e^-^ read noise cameras on the light-sheet microscope. The ∼1.1 μm axial resolution of the light-sheet, while lower than spinning disk confocal, was easily sufficient for Nyquist sampling of 5-6 μm nuclei used to identify each β-cell in 3D space (β-cells themselves are 12- 18 μm). Together these features allowed sampling the islet at >2 Hz, although future studies could be improved by employing a higher numerical aperture objective and a camera with lower read noise and higher quantum efficiency.

Phase and functional network analyses were used to understand the behavior of β-cell subpopulations and how they communicate. Importantly, in past heterogeneity studies, phase and network calculations were assessed over the entire time course^10,15,27,38,39^. Here, we assessed over individual oscillations to compare subpopulation stability over time. One population of cells are those which we termed ‘early phase cells’ and lead the propagating Ca^2+^ wave. These have also been referred to as ‘leader cells’ or ‘pacemaker cells’ and regulate the oscillatory dynamics^12,37,37^. To our surprise, the early phase cells (i.e. the leader cells) were not consistent over time. The late phase cells, located on the opposite end of the islet, showed a similar shift, with over half of the islets showing changes in the wave axis. Consequently, laser ablation of these early phase or late phase cells would be predicted to have little impact on islet function, as suggested previously by electrophysiological studies in which surface β-cells have been voltage-clamped with no impact on β-cell oscillations^19^, or computational studies in which removal of simulated β-cells had little impact on resulting oscillations^15,40^.

Studies have sought to define whether β-cell intrinsic or extrinsic factors determine the oscillations^10,19,35,41^. Because of the shift in Ca^2+^ wave axis between consecutive oscillations, we conclude that β-cell depolarization is dominated by stochastic properties rather than a pre-determined genetic or metabolic profile. Previous experimental and modeling studies have suggested that the Ca^2+^ wave origin corresponds to the glucokinase activity gradient^15,27^. Consistent with this prediction, pharmacologic activation of glucokinase reinforced the islet region of early phase cells and reduced the wave axis change. Pyruvate kinase activation, despite increasing oscillation frequency, had no effect on leader cells, indicating the wave origin is patterned by fuel input. Importantly, there is no evidence that the glucokinase gradient is the result of intentional spatial organization. Rather, in computational studies the glucokinase gradient emerges stochastically due to randomly placed high and low glucokinase- expressing cells, with multiple competing glucokinase gradients determining the degree of wave axis rotation. Our findings suggest that when glucokinase is activated, the strongest gradient is amplified, which is why the Ca^2+^ wave axis is reinforced. Another compelling hypothesis for stochastic behavior, which is not mutually exclusive, is the heterogenous nutrient response of neighboring α-cells influences the excitability of neighboring β-cells via GPCRs^42–44^. The preponderance of α-cells on the periphery of mouse islets, which influence β-cell oscillation frequency^45^, would be expected to disrupt β-cell synchronization on the periphery and stabilize it in the islet center – which is precisely the pattern of network activity we observed. In addition to α-cells, vasculature may also impact islet Ca^2+^ responses^46^, and may induce additional heterogeneity *in vivo*.

Functional network studies of the islet revealed a heterogeneity in β-cell functional connections^14^. A small subpopulation of β-cells, termed ‘hub’ cells, was found to have the highest synchronization to other cells^10^. Optogenetic silencing of hub cells was found to disrupt network activity within that plane, however it should be noted that hub cells are defined as the most highly coordinated cells within a randomly selected plane of the islet. Debates exist over whether the hub cells can maintain electrical control over the whole islet^19,47^. Because our study investigated the 3D β-cell functional network over individual oscillations, our top 10% of highly coordinated cells are not the exact same population as hub cells defined in^10^, however the subpopulations likely overlap. In contrast with leader cells, we found that the highly synchronized hub cells are both spatially and temporally stable. However, in conflict with the description of hub cells as intermingled with other cells throughout the islet^10^, the location of such cells in 3D space is close to the center. This observation could be explained by the peripheral location of α- cells as discussed above for the Ca^2+^ wave behavior.

Previous studies indicated that the intrinsic metabolic activity and thus oscillation profile may play a larger role in driving high synchronization than the strength of gap junction coupling^35,41^. This included experimental 2D measurements but also computational 3D measurements. Nevertheless, we demonstrated here that perturbing glucokinase or pyruvate kinase had little effect on the consistency of the high degree or low degree cells within the 3D network. We further observed that the β-cell network was more regionally consistent than cellularly consistent, indicating a tendency for nearby cells within the islet center to ‘take over’ as high degree cells. The mechanisms underlying this are unclear. One explanation may be that paracrine communication within the islet determines which region of cells will show high or low degree^45^. For example, more peripheral cells that are in contact with nearby δ-cells may show some suppression in their Ca^2+^ dynamics^48^, and thus reduced synchronization. Alternatively, more peripheral cells may show increased stochastic behavior that reduces their relative synchronization. Modulating α/δ-cell inputs to the β-cell in combination with 3D islet imaging will be important to test this in the future. Our study emphasizes that 3D studies are critical to fully assess the consistency and spatial organization of the β-cell network.

## Methods

### Mice

*Ins1-Cre* mice^49^ (Jax 026801) were crossed with *GCaMP6s* mice (Jax 028866), a Cre-dependent Ca^2+^ indicator strain, and *H2B-mCherry* mice, a Cre-dependent nuclear indicator strain^50^. The resulting *Ins1- Cre:ROSA26^GCaMP6s/H2B-mCherry^* mice were genotyped by Transnetyx. Mice were sacrificed by CO_2_ asphyxiation followed by cervical dislocation at 12-15 weeks of age, and islets were isolated and cultured as detailed in^24^. All procedures involving animals were approved by the Institutional Animal Care and Use Committees of the William S. Middleton Memorial Veterans Hospital and followed the NIH Guide for the Care and Use of Laboratory Animals.

### Light-sheet microscope

The stage of a Nikon Ti inverted epifluorescence microscope was replaced with a Mizar TILT M-21N lateral interference tilted excitation light-sheet generator^25^ equipped with an ASI MS-2000 piezo z-stage and Okolab stagetop incubator. The sample chamber consisted of an Ibidi 4-well No. 1.5 glass bottom chamber slide with optically clear sides. Excitation from a Vortran Stradus VeraLase 4-channel (445/488/561/637) single mode fiber-coupled laser and CDRH control box (AVR Optics) was passed through the TILT cylindrical lens to generate a light-sheet with a beam waist of 4.3 μm directly over the objective’s field of view. Similar to a wide-field microscope, the axial (z) resolution of the light-sheet microscope is dictated by the numerical aperture of the objective (∼1.1 μm for our Nikon CFI Apo LWD Lambda S 40XC water immersion objective with a numerical aperture of 1.15). Fluorescence emission was passed through an optical beamsplitter (OptoSplit III, 89 North) and collected by an ORCA-Flash4.0 v3 digital CMOS camera (Hamamatsu C13440-20CU) equipped with a PoCL camera link cable. To achieve high-speed triggered acquisition, the laser and piezo z stage were triggered directly by the camera, which received a single packet of instructions from the NIS-Elements JOBS module via a PCI express NiDAQ card (PCIe-6323, National Instruments). Electronic components (DAQ card, camera, lasers, stage) were linked by a Nikon ‘standard cable’ via a Nikon BNC breakout box; cable assembly is diagrammed in Suppl. Fig. 1. Images were streamed to a Dell computer equipped with an Intel Xeon Silver 4214R CPU, 256 GB RAM, XG5 NVMe SSD, and NVIDIA Quadro Pro 8 GB graphics card and Bitplane Imaris Software (Andor).

### Imaging of β-cell Ca^2+^ and nuclei

Reagents were obtained from Sigma-Aldrich unless indicated otherwise. Islets isolated from *Ins1- Cre:ROSA26^GCaMP6s/H2B-mCherry^* mice were incubated overnight and loaded into an Ibidi µ-slide 4-well No. 1.5 glass bottom chamber slide and maintained by an Okolab stagetop incubator at 37°C. The bath solution contained, in mM: 135 NaCl, 4.8 KCl, 2.5 CaCl2, 1.2 MgCl2, 20 HEPES, 10 glucose, 0.18 glutamine, 0.15 leucine, 0.06 arginine, 0.6 alanine, pH 7.35. Glucokinase activator (50 nM RO 28-1675, Axon), pyruvate kinase activator (10 μM TEPP-46, Calbiochem), and vehicle control (0.1% DMSO) were added as indicated. GCaMP6s (488 nm, 5% power, 50 mW Vortran Stradus Versalase) and H2B- mCherry (561 nm, 20% power, 50 mW) were simultaneously excited and emission was simultaneously collected on a single camera chip using an optical beamsplitter (Optosplit III, 89 North) containing a dichroic mirror (ZT568rdc, Chroma) and emission filters for GCaMP6s (ET525/40, Chroma) and mCherry (ET650/60, Chroma). The exposure time was set to 15 ms in NIS-Elements JOBS, which includes ∼10 ms camera integration time and ∼5 ms stage dwell time. This was sufficiently fast to image intact islets an axial (z) depth of 132 μm at 2.02 Hz (33 z-steps every 4 μm). Raw NIS-Elements ND2 files were imported into Bitplane Imaris analysis software. The location of each cell was marked using H2B-mCherry nuclear signal and a sphere mask was created based on average β-cell nuclear diameter. Nuclear ROIs were mathematically expanded to 9.3 μm to avoid overlapping cells, which was determined by point-scanning confocal imaging^30^. Masks were propagated to all the timepoints and mean Ca^2+^ levels were used to generate single cell traces that were exported from Imaris to Microsoft Excel. Quantitative analyses of the β-cell network and Ca^2+^ wave were performed in MATLAB as described below.

### Identification of β-cell Ca^2+^ Oscillations

To compare β-cell subpopulations over multiple oscillations, we developed a semi-automated oscillation identifier to ensure that the results did not depend on manual identification of oscillation start and end times. First, the approximate time corresponding to the peak of each oscillation was manually identified based on the average islet signal. The time course around each oscillation peak was automatically extracted as seconds before the islet begins depolarization *x*/2 seconds after the islet completes repolarization, where *x* = ¼ oscillation duty cycle. Depolarization and repolarization were then automatically identified using the derivatives of the Ca^2+^ time course and MATLAB’s findpeaks function. All oscillation time courses were manually confirmed. For studies of glycolysis, we ensured that all pre- and post-glycolytic activator treatments had the same number of oscillations. All islets analyzed exhibited slow Ca^2+^ oscillations (period = 6.77 ± 0.36 min).

### Network analysis

Network analysis was conducted as described in^35^, with the caveat that the functional network was recalculated for each oscillation. The correlation threshold was calculated such that the average degree was 7 when averaged over all oscillations^31^.

### Wave analysis

Lagged cross correlation between the normalized Ca^2+^ dynamics of each cell and the islet mean was calculated for each oscillation. Each cell was assigned a cell phase, defined as the time lag that maximized the cross correlation.

### Wave axis

Wave axis was defined as the primary axis between the early (top 10%) and late (bottom 10%) of cells in the Ca^2+^ wave. The primary axis was identified using principal component analysis. To ensure the axis was not confounded by spuriously located cells, cells were not included in the analysis if they were greater than 50 𝜇𝜇m from the center of gravity (calculated using Euclidean distance) of their respective group (early or late phase). Variability of the wave axis over oscillations was defined as the squared Euclidean distance between each wave axis. To compare across islets of differences sizes, wave axis variability was normalized by the maximum variability possible for each islet. This maximum variability was identified by repeating the wave axis calculation 50,000 times for randomly selected early and late phase cells.

### KL-divergence

To calculate consistency over oscillations of the entire islet, network and wave analysis were conducted and cells were ranked for each oscillation (*i*) based on (a) their degree or phase and (b) their Euclidean distance to the center of gravity of the high degree or early phase cells. The probability density functions (𝑝_𝑖_) of these rankings was calculated using the MATLAB normpdf function. The KL-divergence^51^ (𝐷_*KL*_ ) was defined by equation (1), where 𝑛 is the index of each cell, and 𝑖, 𝑗 are indices for each oscillation.

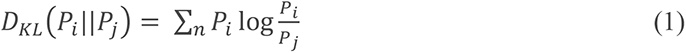

To compare across islets of differences sizes, we normalized the KL divergence by the maximum KL divergence for each islet identified by shuffling the distribution and calculating KL divergence 100 times.

### Comparison of 2D and 3D analyses

Quarter- and half-depth 2D planes were selected from each 3D islet. Cells were included in the 2D plane if their location on the z-axis was within 3 μm of the plane.

### Statistical analysis

Stastistical analysis was conducted using GraphPad PRISM 9.0 software. Significance was tested by first testing normality using Anderson-Darling and Kolmogorov-Smirnov normality tests and then using paired Wilcox tests, Student’s two-tailed t-tests or ANOVA as indicated. *P* < 0.05 was considered significant and errors signify ±SEM.

## Data and code availability

All data analyzed during this study will be included in this published article and its supplementary information files. All code is publicly available at https://github.com/jenniferkbriggs/Lightsheet_BetaCell_Identity

## Acknowledgements

We thank Barak Blum at the University of Wisconsin-Madison for providing *Rosa26^H2B-mCherry^* mice and the University of Wisconsin Optical Imaging Core for use of the spinning disk confocal. The Merrins laboratory gratefully acknowledges support from the NIH/NIDDK (R01DK113103 and R01DK127637 to M.J.M., and R01DK106412 to R.K.P.B.) and the United States Department of Veterans Affairs Biomedical Laboratory Research and Development Service (I01BX005113 to M.J.M.). The Benninger laboratory gratefully acknowledges support from the NIH/NIDDK (R01DK106412, R01DK102950, R01DK140904 to R.K.P.B.) and the University of Colorado Diabetes Research center (P30 DK116073). Jennifer K Briggs acknowledges support from NSF GRFP (DGE-1938058_Briggs).

## Author Contributions

E.J. constructed the microscope and performed islet imaging experiments. J.K.B. and E.J. analyzed the data and created the figures. E.J. and M.J.M. drafted the paper and all authors edited the paper. M.J.M and R.K.P.B. provided resources.

## Competing interests

The authors declare no competing interests.

**Correspondence** and requests for materials should be addressed to Matthew J. Merrins and Richard K.P. Benninger.

---

**Suppl. Fig. 1.**
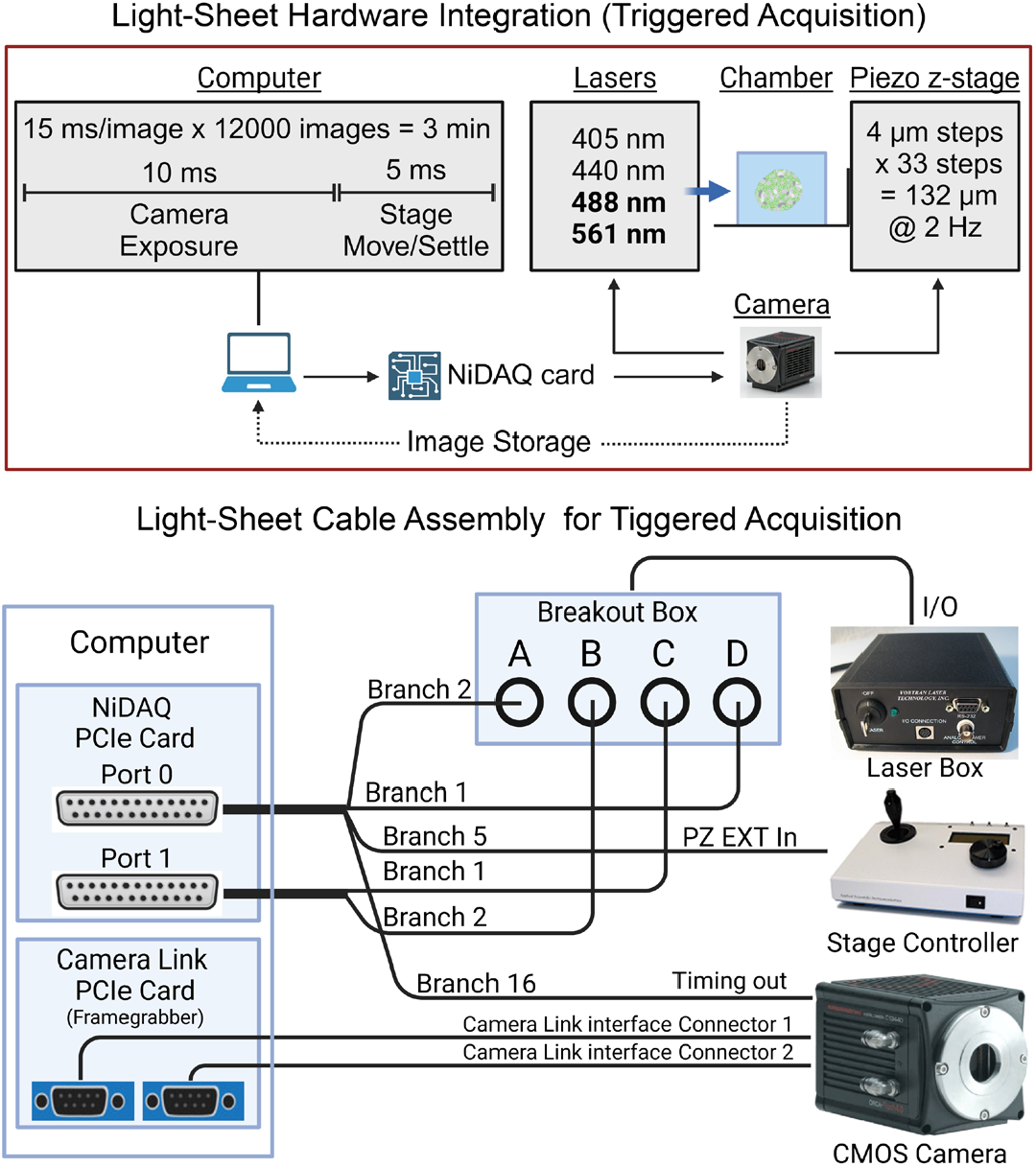
Hardware wiring diagram of the light-sheet microscope. Hardware integration (top panel) for camera-triggered activation of the excitation lasers and piezo z-stage that limits communication to a single instruction from the computer every 3 minutes. Wiring diagram (bottom panel): two Nikon ‘standard cables’ connect to the NiDAQ card installed in the computer. These two cables link to the laser control box, stage controller and camera. The images captured are received by the computer through a camera link PCIe card. and

**Suppl. Fig. 2.**
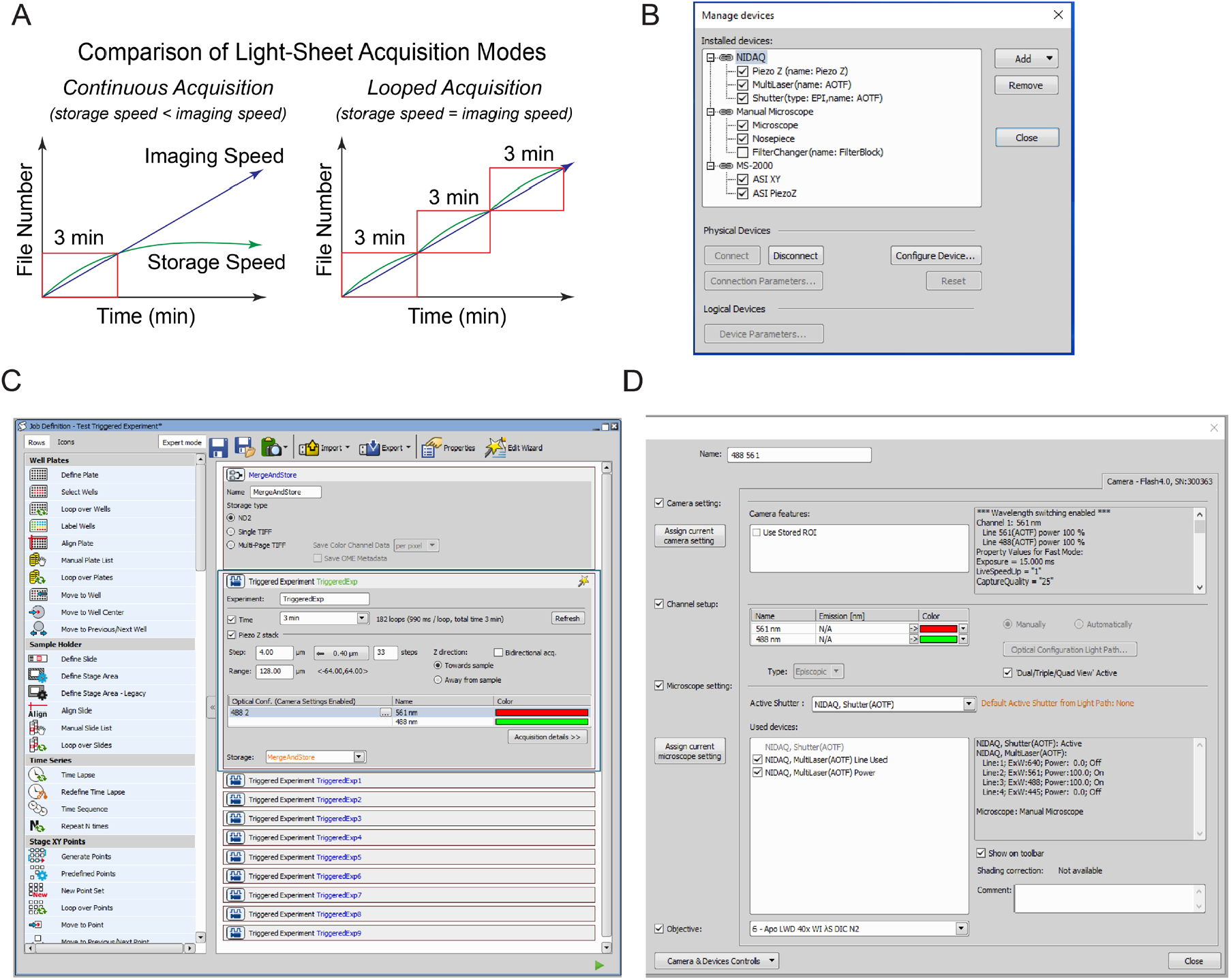
NIS-Elements software configuration. (A) Schematic for comparing continuous acquisition and looped acquisition. The red box indicates the 3-minute window where storage speed is higher than imaging speed. (B) Devices linked to NIS-Elements. (C) NIS-Elements JOBS module configured for looped acquisition. (D) Optical configuration for simultaneous GCaMP6s/H2B-mCherry excitation.

**Suppl. Fig. 3.**
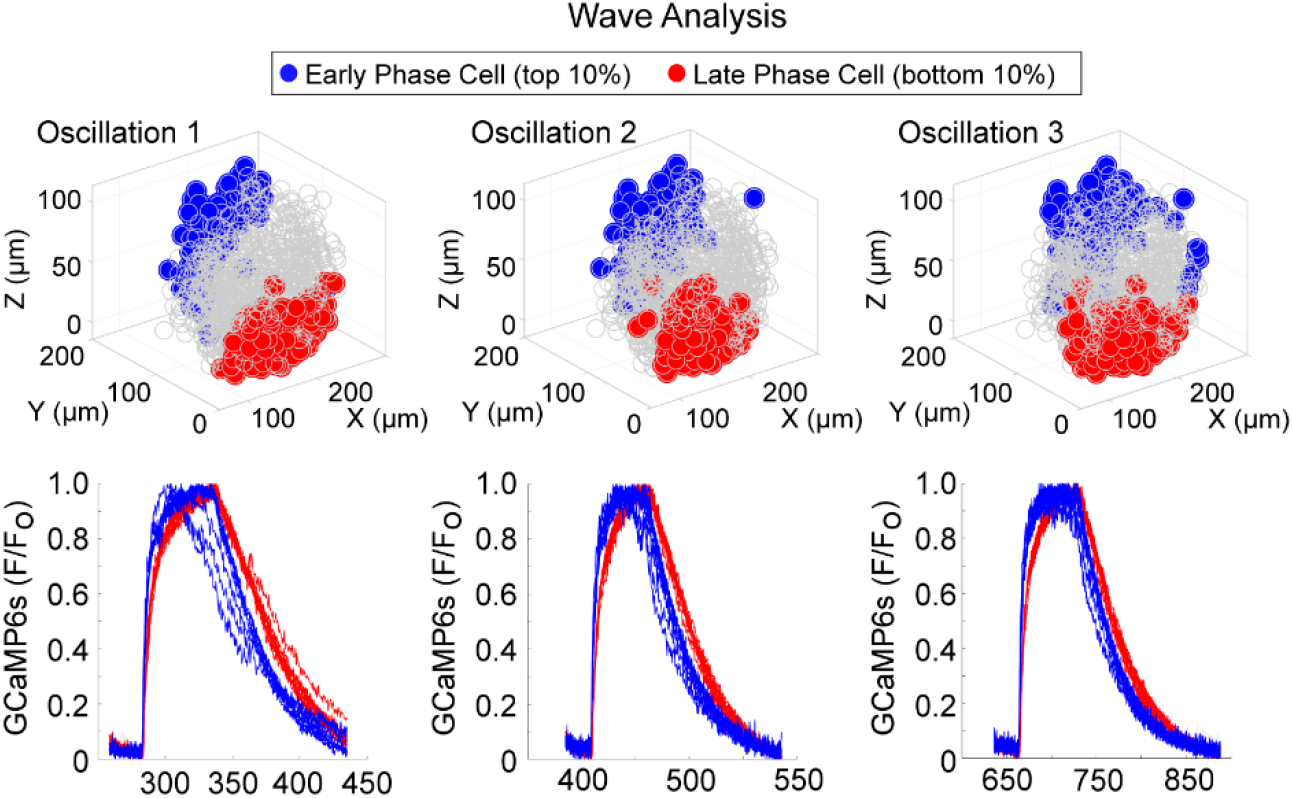
Location of early and late phase cells in an islet with stable wave axis. 3D representation of the islet showing the location of early phase cells (blue) and late phase cells (red) over three consecutive oscillations (top panel) and their corresponding Ca^2+^ traces (bottom panel).

**Suppl. Fig. 4.**
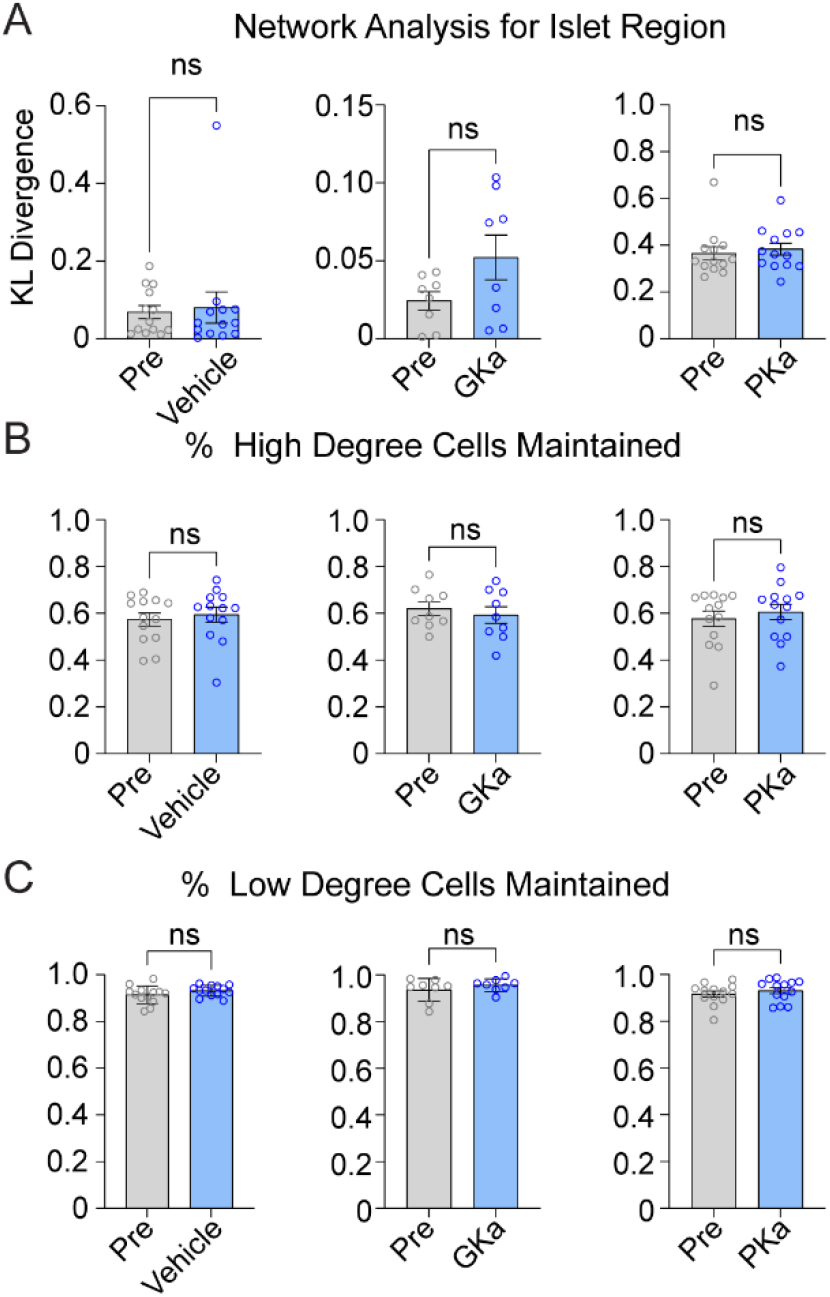
Effect of glycolytic activators on the β-cell network. (A-C) Effect of vehicle, glucokinase activator (GKa), and pyruvate kinase activator (PKa) on regional consistency of the β-cell network (A), high degree cell retention (B), and low degree cells retention (C). Data are displayed as mean ± SEM.

**Suppl. Fig. 5.**
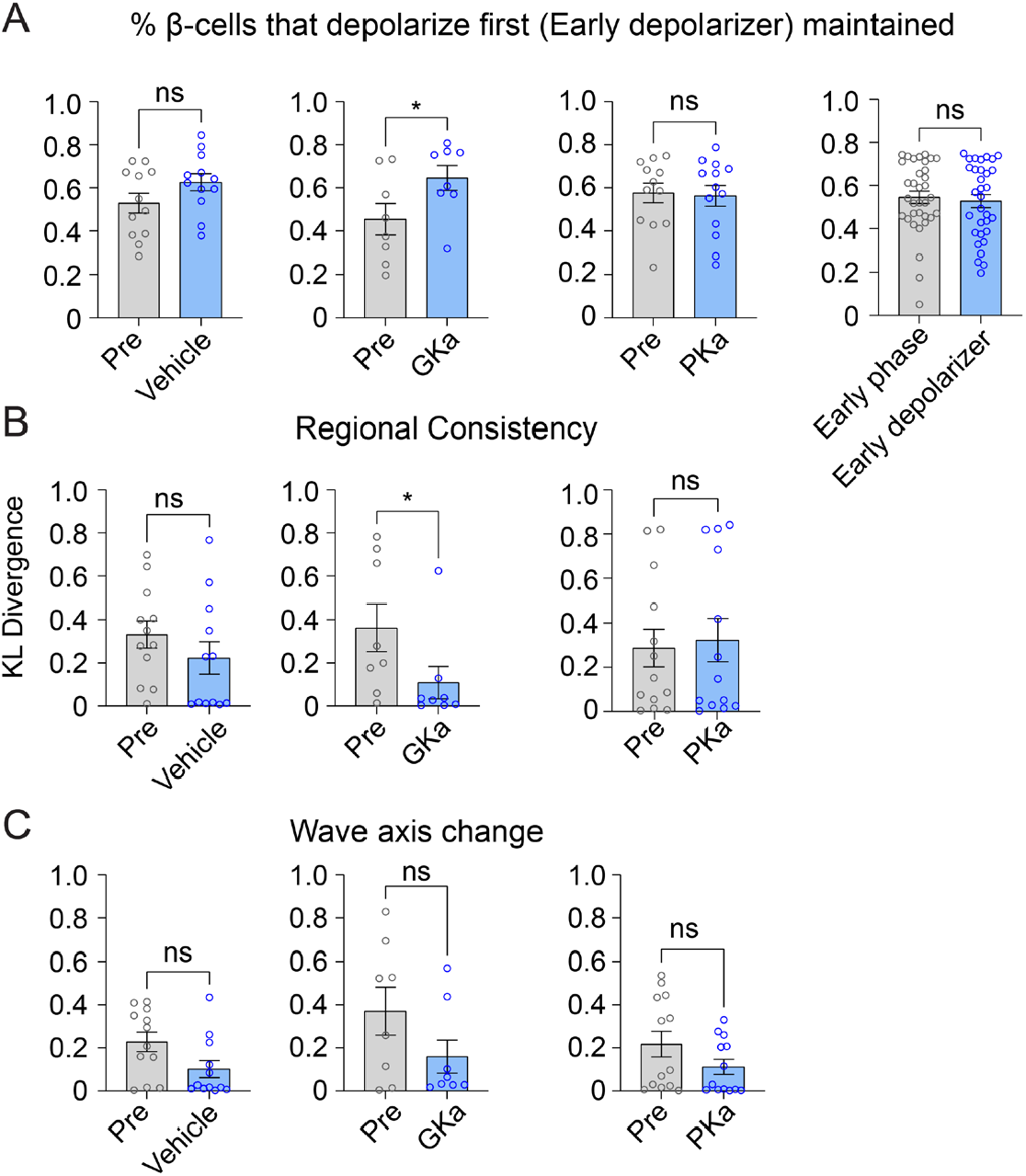
Effect of glycolytic activators on the β-cells that depolarize first. (A-C) Effect of vehicle, glucokinase activator (GKa), and pyruvate kinase activator (PKa) on the retention (A), regional consistency (B), and wave axis change (C) of the β-cells that depolarize first (Early depolarizer). The retention of β-cells that depolarize first (Early depolarizer) is similar to cells that depolarize and repolarize first (Early phase). Data are displayed as mean ± SEM.

